# Systematic enhancement of protein crystallization efficiency by bulk lysine-to-arginine (KR) substitution

**DOI:** 10.1101/2023.06.03.543563

**Authors:** Nooriel E. Banayan, Blaine J. Loughlin, Shikha Singh, Farhad Forouhar, Guanqi Lu, Kam-Ho Wong, Matthew Neky, Henry S. Hunt, Larry B. Bateman, Angel Tamez, Samuel K. Handelman, W. Nicholson Price, John F. Hunt

## Abstract

Structural genomics consortia established that protein crystallization is the primary obstacle to structure determination using x-ray crystallography. We previously demonstrated that crystallization propensity is systematically related to primary sequence, and we subsequently performed computational analyses showing that arginine is the most overrepresented amino acid in crystal-packing interfaces in the Protein Data Bank. Given the similar physicochemical characteristics of arginine and lysine, we hypothesized that multiple lysine-to-arginine (KR) substitutions should improve crystallization. To test this hypothesis, we developed software that ranks lysine sites in a target protein based on the redundancy-corrected KR substitution frequency in homologs. We demonstrate that three unrelated single-domain proteins can tolerate 5-11 KR substitutions with at most minor destabilization and that these substitutions consistently enhance crystallization propensity. This approach rapidly produced a 1.9 Å crystal structure of a human protein domain refractory to crystallization with its native sequence. Structures from bulk-KR-substituted domains show the engineered arginine residues frequently make high-quality hydrogen-bonds across crystal-packing interfaces. We thus demonstrate that bulk KR substitution represents a rational and efficient method for probabilistic engineering of protein surface properties to improve protein crystallization.

---

More than 50 years after the solution of the first protein crystal structure ^1–3^, protein crystallization remains a hit-or-miss proposition. Synergistic developments in crystallographic methods^4–9^, synchrotron beamlines^10–13^, and high-speed computing have made structure solution and refinement routine, even for massive complexes, as long as high-quality crystals are available. However, there has been comparatively little progress in improving methods for protein crystallization. Structural genomics consortia have systematically confirmed that most naturally occurring proteins do not readily yield high-quality crystals suitable for x-ray structure determination and that crystallization is the major obstacle to the determination of protein structures using diffraction methods ^14–16^. While numerous methods have been developed that have some efficacy in improving protein crystallization properties ^17–27^, none work with sufficiently high efficiency to have been applied with significant frequency by practicing crystallographers. We therefore set out to develop efficient methods for rational engineering of protein surface properties to improve crystallization propensity. The first phase of our research identified a large number of local primary sequence patterns, which we called crystallization epitopes, that are strongly overrepresented in crystal-packing interfaces ^28^. We demonstrated that introducing these epitopes individually into proteins generally increases their crystallization propensity and that introducing multiple such epitopes progressively increases crystallization propensity. The cumulative nature of the observed improvements suggested that multiple simultaneous mutations could potentially produce definitive improvements in crystallization propensity in a single protein construct based on large-scale probabilistic engineering of protein surface properties. We herein present an efficient method to achieve this goal while preserving protein stability, solubility, and function.

Our efforts to develop rational methods to improve protein crystallization properties are grounded in sequence and structural analyses of historical crystallization results and associated thermodynamics studies. Our published analyses of large-scale experimental studies showed that the surface properties of proteins, and particularly the entropy of the exposed sidechains, are a major determinant of protein crystallization propensity ^16^. These studies demonstrated that overall thermodynamic stability is not a major determinant of protein crystallization propensity. They identified a number of primary sequence properties that correlate with successful crystal structure determination, including strong anticorrelations with predicted backbone disorder and surface sidechain entropy and weak positive correlations the fractional content of several individual amino acids ^16^. In follow-up studies, we analyzed 87,683 crystal structures from the Protein Data Bank (PDB) and identified contiguous amino acid patterns strongly overrepresented in crystal packing interfaces ^28^. This analysis also generated data on the relative overrepresentation of individual amino acids in crystal-packing interfaces segregated by secondary structure (**Fig. 1**), and these data suggested the streamlined approach reported in this paper that enhances protein crystallization propensity based on multiple simultaneous surface mutations.

**Figure 1.**
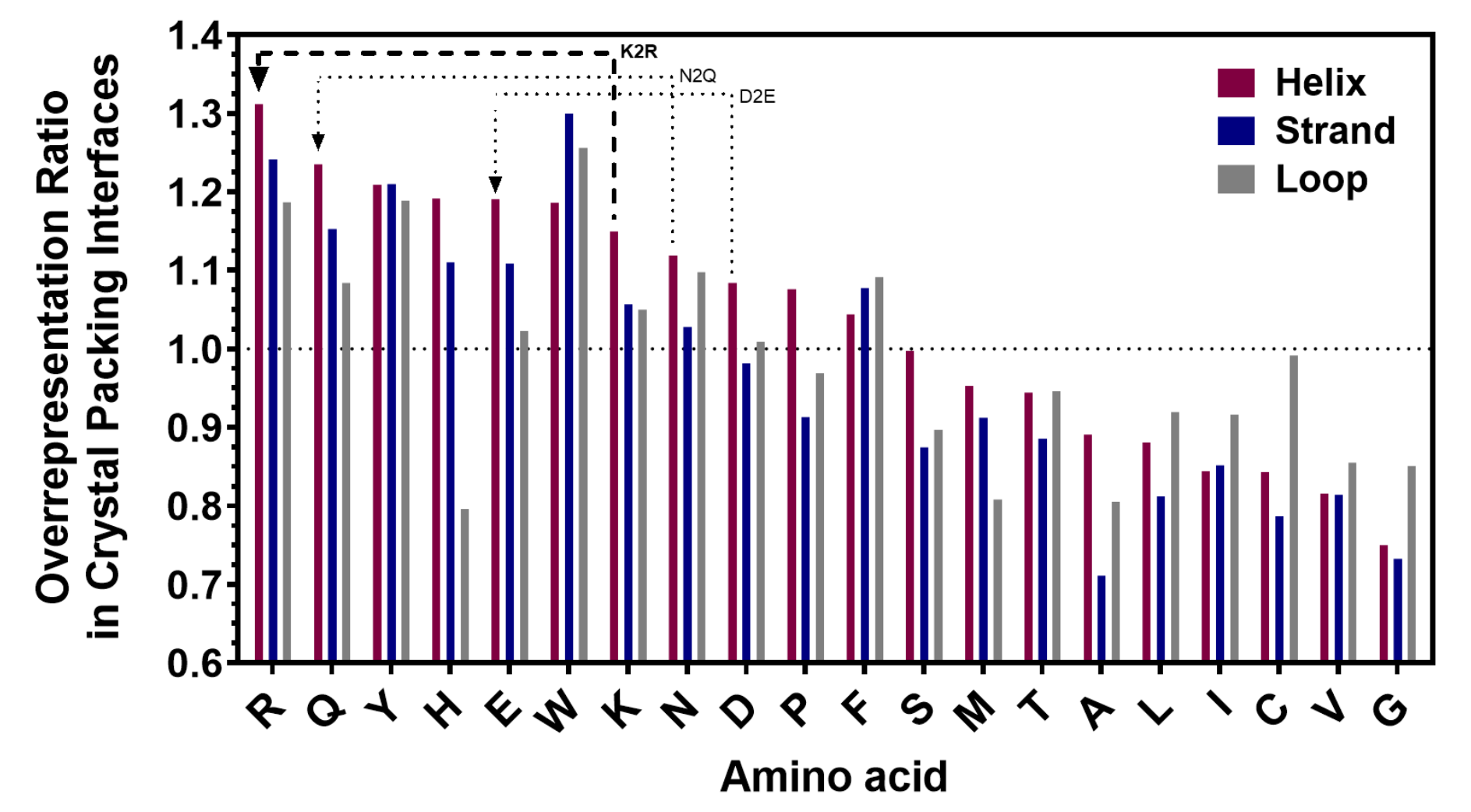
Overrepresentation ratios ^28^ of amino-acids in crystal-packing interfaces compared to overall surface composition in 87,684 crystal structures deposited in the Protein Data Bank (PDB). The red circles/arrows schematize the gain in packing probability produced by K-to-R mutations. The overrepresentation ratios are segregated by protein 2° structure as assessed by DSSP ^82^, and the amino acids are ordered in decreasing order of overrepresentation ratio in α- helical secondary structure.

Our computational analysis of crystal-packing interactions in the PDB showed a substantially higher probability for arginine to mediate inter-molecular packing contacts than lysine (**Fig. 1**), consistent with our expectations based on earlier analyses of correlations between primary sequence features and protein crystallization propensity ^16^. The observation that arginine mediates crystal-packing contacts more frequently than lysine is particularly notable because the entropy of the arginine sidechain is generally estimated to be somewhat higher than that of lysine ^29–32^, and higher surface sidechain entropy opposes crystallization ^16, 17, 22–24^. Therefore, the more frequent occurrence of arginine compared to lysine in crystal-packing contacts suggests that the guanidino group on arginine is substantially “stickier”, in terms of intermolecular interaction free energy, than the primary amine on lysine. This inference is consistent with the well-known properties of the guanidinium ion as a protein denaturant ^33–35^, a property that is not shared by primary amines. Therefore, we hypothesized that introducing multiple arginine-to-lysine (KR) substitutions in a protein would enhance crystallization propensity and that, given the very similar physicochemical properties of arginine and lysine in terms of size and polarity, multiple simultaneous substitutions would be tolerated without significantly impairing thermodynamic stability.

We herein report the results of biophysical studies that support the validity of this hypothesis. We developed a computer program that automates selection of sites for KR mutagenesis based on the frequency of such substitutions in naturally occurring homologs, which should avoid sites where lysine is critical for function or structural stability. We furthermore characterized the effects of introducing multiple simultaneous KR mutations on the thermodynamic stability, solubility and crystallization propensity of three unrelated test proteins, one of which crystallizes readily and two of which are recalcitrant to crystallization with their native sequences. These studies demonstrate that introducing multiple KR mutations into a protein, which we call Bulk KR substitution, is a simple and effective method to improve crystallization propensity. Physicochemical analyses have thus guided the development of an efficient method for large-scale probabilistic engineering of protein surface properties to improve crystallization, which was historically considered a stochastic phenomenon refractory to rational experimental manipulation.

## Results

### KR mutation site-selection algorithm and software

Sites for Bulk KR substitution are ranked based on the frequency of these substitutions in naturally evolved sequences in a phylogenetic alignment. This procedure is fully automated in Python code available at Github (https://github.com/huntmolecularbiophysicslab/pxengineering) that can be run interactively via a webserver that can be accessed at http://www.pxengineering.org. The algorithm implemented by the program ranks sites based on a redundancy-compensated estimate (explained below) of the frequency of KR substitutions in homologous sequences, which are divided into mutually exclusive bins with progressively lower levels of overall percent identity relative to the target sequence. The first bin includes sequences with greater than or equal to 90% identity and less than 99% identity (to avoid mutant variants of the target sequence), and subsequent mutually exclusive bins reduce the lower identity level in 10% steps down to a minimum of 30%. The algorithm steps through these bins in order from highest to lowest percent identity, selecting sites in each bin in inverse order of their redundancy-compensated count of lysine-to-arginine substitutions down to a minimum user-adjustable threshold count. This threshold is imposed to avoid selecting a site based on an arginine substitution in a single sequence that could potentially be inaccurate or in a small number of very closely related sequences that could potentially share a function-impairing or stability-impairing mutation; it defaults to a value of 1.1, which ensures observation of a lysine-to-arginine substitution in at least two sequences with no more than ∼93% identity to one another.

The software provides graphical displays of summary parameters characterizing the amino acid distribution in the homologs in each of the percent-identity bins at every lysine site in the target sequence (**Fig. 2**), as well as a graphical display of the overall sequence diversity in each of the bins (**Fig. ED1**). The displayed summary parameters are the Shannon entropy of the amino-acid frequency distribution, the frequency of all residues other than lysine, the arginine-to-lysine ratio, the total count of sequences with an arginine residue at the site, and two different estimates of that count after compensation for redundancy between those sequences. Both redundancy-compensation calculations use the same heuristic estimate of the degree of mutational resampling between pairs of sequences, which is described in the **Methods** section along with explanations of the details of the two algorithms. In brief, the first redundancy-reduced count evaluates all sequences using a calculation that has rigorously correct behavior in the cases of full redundancy and full independence between the sequences but is otherwise approximate. The second count provides a rigorous probabilistic estimate of the number of independent observations in the seven most remotely related sequence pairs in the bin having arginine at that site. Extending this calculation to more sequences is computationally prohibitive, but the estimate based on a limited set of the most diverged homologs provides a highly effective method to ensure that multiple independently determined protein sequences have an arginine residue at the lysine site in the target protein, which is the essential goal of the redundancy-compensation calculations. This second calculation is used for the automated site-ranking algorithm described above.

**Figure 2.**
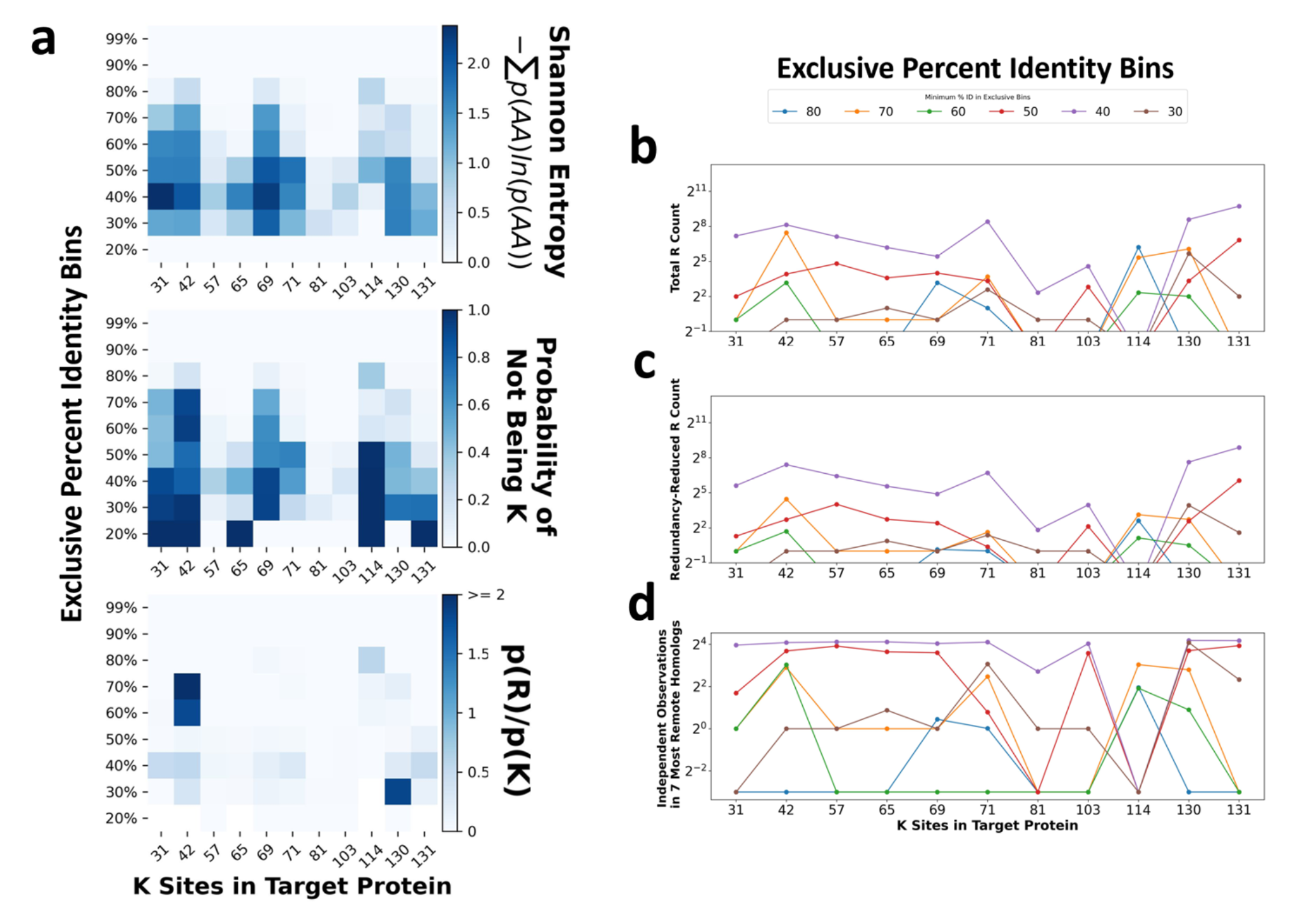
Representative output from the Bulk-R webserver analyzing lysine-to-arginine substitution patterns in homologous proteins. All sequence analyses are conducted on sets of proteins spanning 10% ranges (bins) of sequence identity relative to the target protein, with the exception of the highest identity bin which spans 90-99% identity to avoid mutant protein sequences. **(a)** The top graph shows the Shannon entropy, the middle graph shows the fraction of residues other than lysine, and the bottom graph shows the ratio of arginine/lysine residues at each lysine site in the native sequence in the indicated sequence identity bin. **(b,c)** The total (panel **b**) and heuristically redundancy-reduced (panel **c**) counts of arginine residues observed at each lysine site in the native sequence in the indicated sequence identity bin. **(d)** The expectation value for the number of independent observations of arginine in up to the seven most diverged sequence pairs in the bin that have arginine at the indicated site. The graph in panel **c** displays the results of the first redundancy-correction calculation described in the text, while the graph in panel **d** displays the results of the second.

The program additionally provides a ranking of sites for introducing aspartate-to-glutamate and asparagine-to-glutamine mutations together with a record of which of those sites have potential ionic interaction or H-bonding partners in the target sequence that would tend to reduce the entropy of the longer sidechains in beta-sheet (i±2) or α-helical (i±3,i±4) secondary structures ^36–38^. (The rationale behind this approach is described in the ***Discussion*** section below.) Lysine, arginine, and histidine are considered potential ionic interaction partners for glutamate and H-bonding partners for glutamine, while asparagine, glutamine, serine, and threonine are considered potential H-bonding partners for both with the addition of aspartate and glutamate for glutamine.

### Test protein selection and expression

We chose to test the Bulk KR substitution approach using three proteins with different crystallization properties. The hPDIa domain is a human drug target ^39, 40^ that represents the first of four domains in the endoplasmic-reticulum-resident human Protein Disulfide Isomerase (hPDI) protein. The hPDIa domain had never successfully been crystallized on its own, but its structure was known from a relatively low-resolution crystal structure of a much longer multi-domain construct containing hPDIa ^41^, which enables evaluation of the impact of Bulk KR substitutions on its structure, as reported below. *E. coli* RNaseH is difficult to crystallize in the absence of ligands stabilizing active site structure but has had its crystal structure determined by groups studying its enzymological mechanism and folding ^42–47^. MA_2137, an S-adenosyl-methione-dependent RNA methyltransferase from *Methanosarcina acetivorans*, crystallizes well in the presence of S-adenosyl-homocysteine (SAH), the product of the methyltransferase reaction that it catalyzes. We included this last protein because we previously demonstrated that increasing hit count in high-throughput crystallization screening is strongly correlated with the probability of successful crystal-structure determination ^16^, which implies quantification of hit count for a protein that crystallizes relatively easily is an effective assay for crystallization propensity. KR mutations were introduced into the D65R mutant of MA_2137 because we had previously demonstrated that this single mutation improves crystallization of this protein, and we wanted to determine whether bulk KR substitutions could improve it even further.

We introduced 2 to 13 KR mutations into these proteins (**Table 1**), and we first examined the expression and solubility levels of the full set of mutant constructs when expressed from a pET plasmid using T7 RNA polymerase in *E. coli*, which yields high-level expression of the three parental proteins in the form of efficiently purified monodisperse monomers. The largest number of KR mutations tested preserved high-yield protein production in a monodisperse state for both hPDIa and MA_2137-D65R (*i.e.*, the hPDIa-9KR and MA_2137-D65R-11KR constructs). The RNaseH-2KR and RNaseH-5KR constructs similarly preserved high-yield protein production in a monodisperse state. However, the RNaseH-7KR construct yielded polydisperse protein that co-purified with the Hsp33 molecular chaperone protein ^48, 49^, while the RNaseH-11KR was completely insoluble even though it expressed at a high level (data not shown). The stability studies presented in the next section confirm earlier research ^50, 51^ showing that RNaseH has a low thermal melting temperature (T_m_) of ∼45 °C, making it marginally stable, which likely explains its tolerance for fewer KR mutations than the other target proteins.

**Table 1.** Summary of expression, stability, and crystallization results for Bulk-KR mutant proteins.*

| Target Protein | Construct | KR mutation sites | Expression | Solubility | T<sub>m</sub> (°C) | ΔH<sub>vH</sub> (kcal/mol) | # hits at 4 weeks | Crystal structure resolution (Å) |
| --- | --- | --- | --- | --- | --- | --- | --- | --- |
| hPDIa<br>120 amino acids | WT | – | +++ | +++ | 59.9, 67.9 | 204 ± 36, 131 ± 9 | 0 | n/a |
|  | 2KR | 42, 114 | +++ | +++ | 58.9, 68.6 | 201 ± 40, 117 ± 5 | n/a | n/a |
|  | 5KR | + 130, 69, 71 | +++ | +++ | 48.2, 65.7 | 47.7 ± 2, 125 ± 3 | n/a | n/a |
|  | 7KR | + 31, 131 | +++ | +++ | 49.5, 64.1 | 89.8 ± 5, 156 ± 5 | n/a | n/a |
|  | 9KR | + 57, 65 | +++ | +++ | 55.4, 60.4 | 163 ± 13, 161 ± 3 | 9 | 1.89 Å |
| MA_2137<br>194 amino acids | WT | – | +++ | +++ | 69.0 | 283 ± 13.5 | 75 | n/a |
|  | D65R | – | +++ | +++ | 68.7 | 250 ± 4.60 | 126 | 1.60 Å |
|  | D65R-3KR | 126, 129, 194 | +++ | +++ | 67.1 | 194 ± 3.30 | n/a | n/a |
|  | D65R-5KR | + 52 71 | +++ | +++ | 67.4 | 153 ± 7.10 | n/a | n/a |
|  | D65R-7KR | + 172, 133 | +++ | +++ | 65.5 | 193 ± 3.90 | n/a | n/a |
|  | D65R-11KR | + 8, 64, 142, 155 | +++ | +++ | 63.4 | 150 ± 2.60 | 238 | 1.91 Å |
| RNaseH<br>166 amino acids | WT | – | +++ | +++ | 45.2 | 135 ± 2.20 | 0 | n/a |
|  | 2KR | 31, 90 | +++ | +++ | n/a | n/a | n/a | n/a |
|  | 5KR | + 66, 89, 35 | +++ | +++ | 45.5 | 128 ± 2.60 | 0 | n/a |
|  | 7KR | + 107, 124 | +++ | + | n/a | n/a | n/a | n/a |
|  | 11KR | + 111, 62, 88, 101 | +++ | – | n/a | n/a | n/a | n/a |

\* The constructs harboring increasing numbers of KR mutations also include all of those in the constructs of the same protein harboring fewer KR mutations. The code “n/a” stands for not applicable or not attempted.

### KR mutations are generally only minimally destabilizing

The thermal stabilities of all the successfully purified bulk KR construct were characterized using circular dichroism spectroscopy. These assays show a variable but generally very small degree of destabilization by KR mutations (**Fig. 3** and **Table 1**). RNase-5KR shows an approximately unaltered T_m_ compared to the wild-type protein, demonstrating that KR mutations can have a completely neutral effect on stability. PDIa-9KR shows an ∼8° reduction compared to the 68 °C T_m_ of the wild-type domain, while MA_2137-D65R-11KR shows an ∼6° reduction compared to the 69 °C T_m_ of the parental protein. Considering the entire set of mutant proteins in our study that could be purified, which includes 25 different KR mutations (**Table 1**), there is on average a 0.54 +/- 0.30° reduction in T_m_ per KR mutation. Therefore, KR mutations are generally very well tolerated, although large sets of mutations tend to produce modest reductions in protein stability ^52^ that can reduce soluble protein yield *in vivo* when the stability of the wild-type (WT) protein is relatively low.

**Figure 3.**
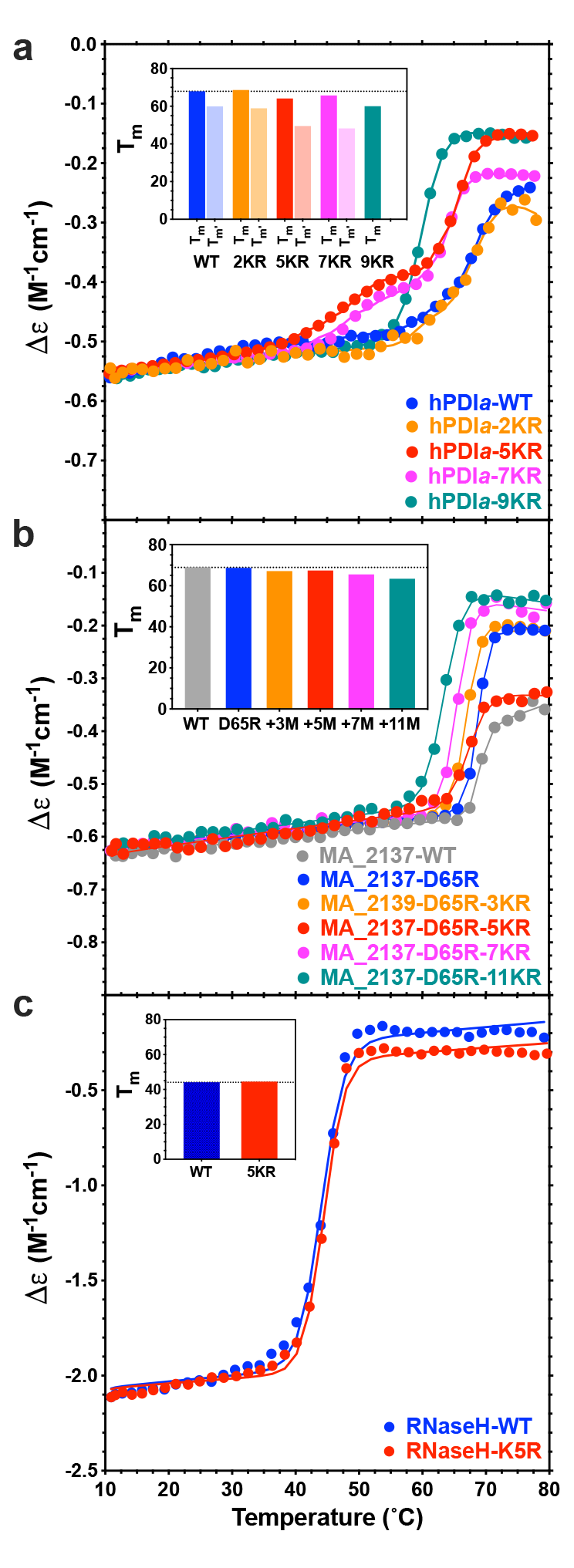
Thermodynamic stability of bulk-KR mutants characterized using thermal denaturation experiments monitored by circular dichroism (CD) spectroscopy. Experiments were conducted using protein samples at 2 mg/ml scanned at a rate of 3 °/min in a buffer containing 100 mM NaCl, 10 mM Tris-Cl, pH 7.5, with the addition of 1 mM SAH for the MA_2317 constructs. The suppression of the low-temperature transition in the hPDIa-9KR construct could be attributable to an intra-helical salt bridge between the sidechains of residues E62 and R65. The latter residue is one of the KR mutations in this construct not shared by the hPDIa-7KR construct, and the sidechain of residue K65 does not make any H-bonds at all in the crystal structure of the multidomain construct (PDB id 4EKZ). Therefore, the sidechain salt-bridge produced by the K65R mutation could potentially stabilize local structure in the hPDIa domain.

### Bulk KR mutations enhance crystallization propensity and yield strongly diffracting crystals

The purified protein constructs harboring the largest number of KR mutations (*i.e.*, PDIa-9KR and MA_2137-D65R-11KR) along with matched controls were screened for crystallization at the National Crystallization Center at the Hauptman-Woodward Institute (HWI) using their automated, high-throughput 1536-condition screen. This well-documented ^53–58^ microbatch-under-oil screen was employed for initial crystallization screening by the Northeast Structural Genomics Consortium^59–62^ (www.nesg.org), which used it to generate 664 crystal structures deposited in the PDB. Neither the WT or 5KR construct of RNaseH yielded any crystallization hits in a screen intentionally conducted without any ligands stabilizing active site structure in order to provide the most exacting test of protein crystallization propensity; the lack of success for this protein was potentially influenced by the high 15 mg/ml protein concentration used for screening, which produced pervasive amorphous precipitation in the screen at the earliest observation times. However, the hPDIa-9KR and MA_2137-D65R-11KR constructs both yielded significantly more crystallization hits than the control proteins. MA_2137-D65R-11KR yielded hits under twice as many conditions as the MA_2137-D65R control protein, while hPDIa-9KR yielded 9 high quality hits compared to no hits at all for the WT construct (**Fig. 4** and **Table 1**). A small number of hit conditions for each protein were chosen for optimization, which very rapidly yielded 1.9 Å structures for both Bulk KR constructs based on a single session of remote synchrotron diffraction screening and data collection (**Figs. 5 & ED2** and **Table ED1**). Therefore, for both target proteins, crystallization screening only had to be conducted on the soluble construct harboring the largest number of Bulk KR mutations in order to rapidly obtain high-quality crystal structures.

**Figure 4.**
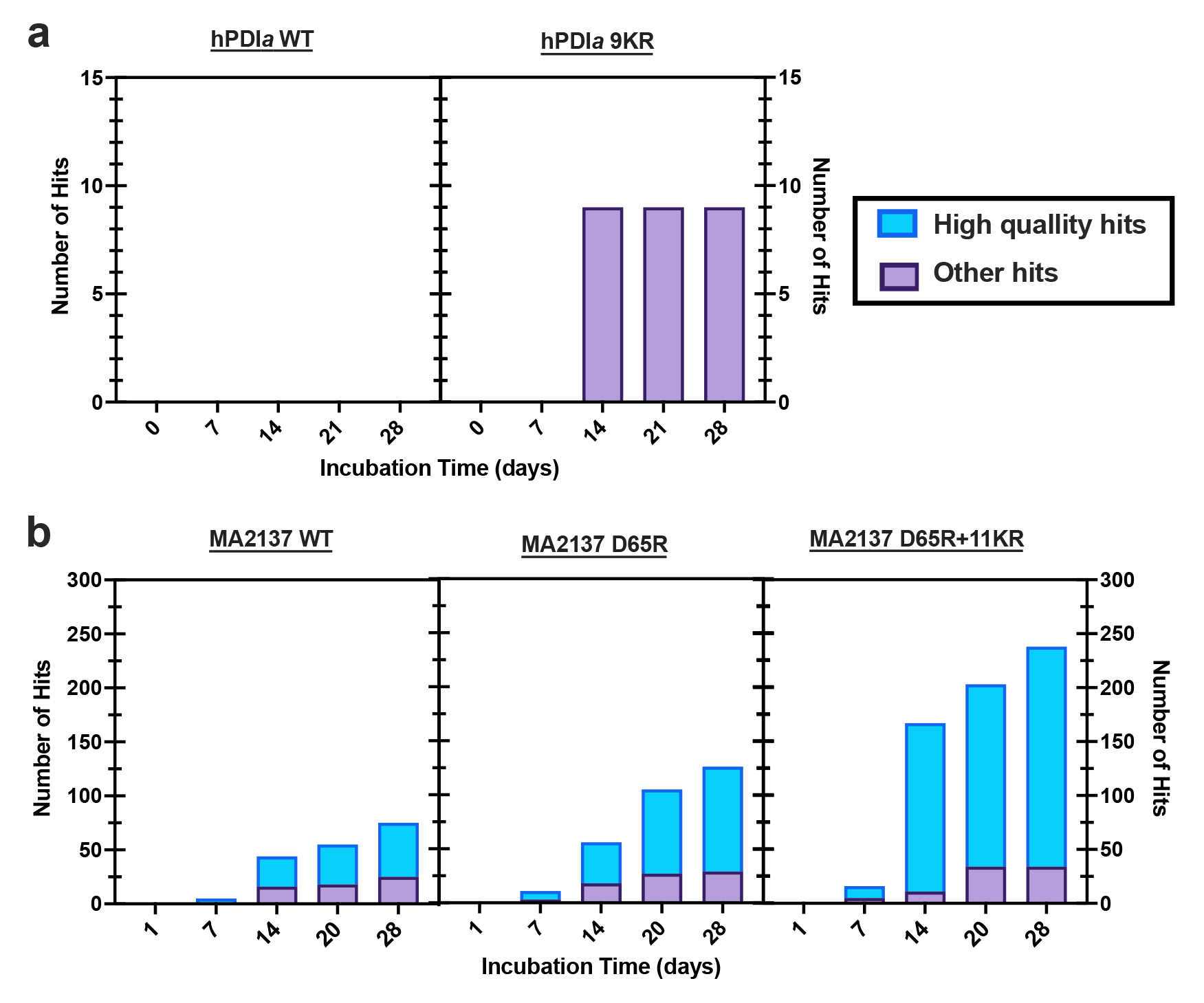
Crystallization hit counts from the 1536-condition high-throughput automated microbatch-under-oil crystallization screen at the National Crystallization Center at the Hauptman-Woodward Institute (HWI). These screens were conducted during the summer of 2021 using generation 19 of the HWI crystallization cocktail collection. The proteins were at ∼15 ml/ml in the stock solutions used for screening, which contained 100 NaCl, 10 mM DTT, 10 mM Tris-Cl, pH 7.5. The screening plates are maintained at 4 °C for the first week and then moved to 25 °C for the remainder of the screening period ^55–57, 83^.

**Figure 5.**
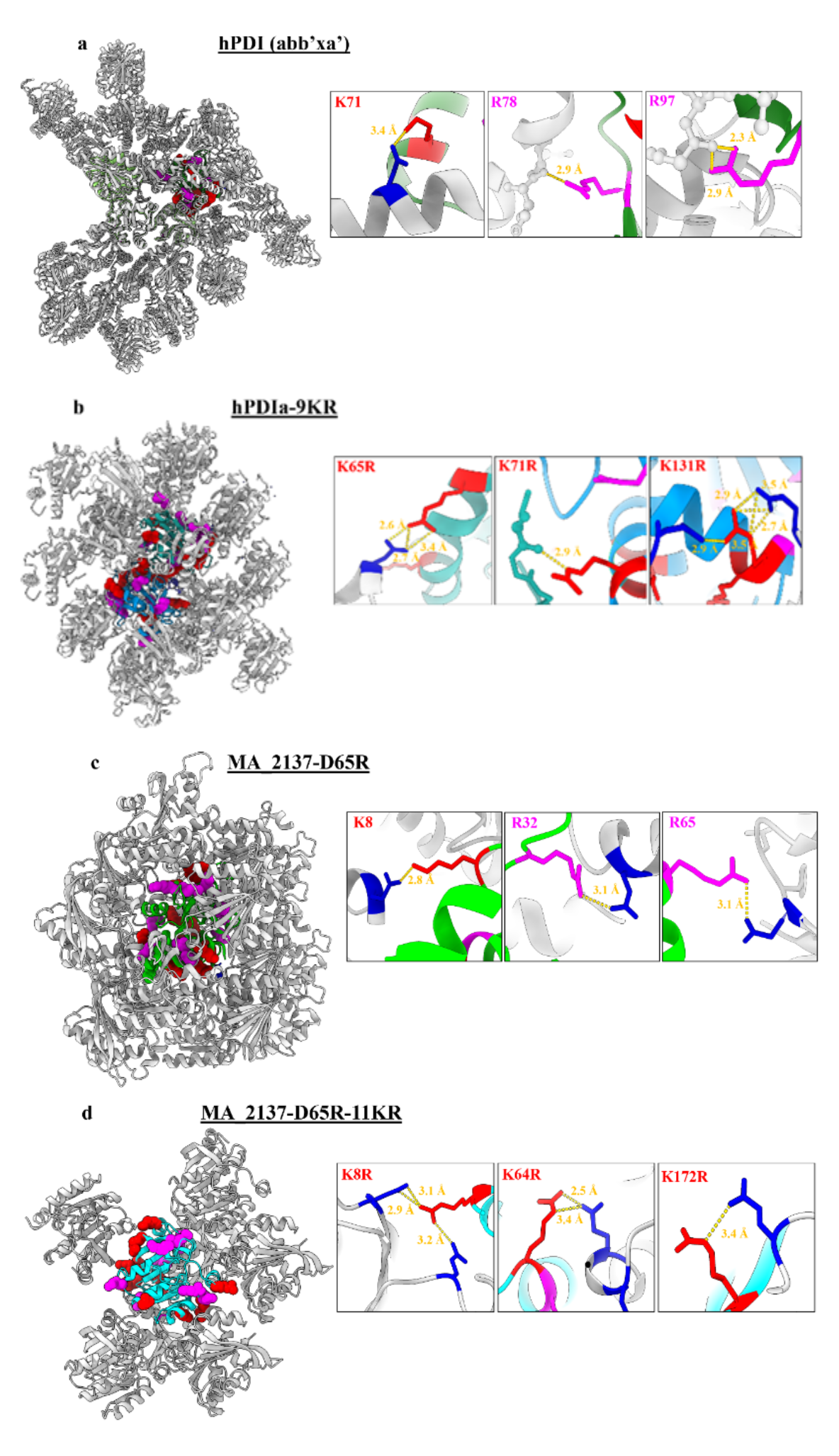
Crystal-packing in structures from proteins containing bulk-KR mutations. The protein backbone is shown in ribbon representation, colored gray for symmetry mates and shades of green, blue, cyan, and teal for the subunits modeled in the asymmetric unit of each crystal structure. The sidechains of the native arginines (magenta) and residues mutated from lysine-to-arginine (red) are shown in space-filling representation on the left and in stick representation in the zoomed-in views of local packing interactions in the boxes to the right. Asp, asn, glu, and gln residues making H-bonds to the illustrated lysine and arginine residues in the boxes are shown in blue stick representation, while the backbone atoms making H-bonds to those residues are shown in ball-and-stick representation.

### Bulk KR mutations frequently make H-bonds in crystal-packing interfaces without perturbing protein structure

The 1.9 Å crystal structures of our Bulk KR substitution (**Fig. 5** and **Table ED1**) constructs show 0.32-0.33 Å root-mean-square deviations for their backbone Cα atoms compared to the references structures (**Fig. ED2**) (*i.e.*, the much larger multidomain hPDI(abb’xa’) construct for hPDIa because the isolated domain has never successfully been crystallized before and the parental MA_2137-D65R construct for MA_2137-D65R-11KR). The observed deviations are close to the expected coordinate error in well-refined crystal structures in the operative resolution range ^63^, indicating our Bulk KR substitution method does not significantly perturb protein conformation for either of our targets. Detailed analyses of the intermolecular interactions in our crystal structures demonstrates that the engineered arginine sidechains make extensive crystal packing contacts, substantially exceeding the number of van der Waals contacts and especially H-bonds made by the native arginine sidechains in the same constructs and greatly exceeding the number of both kinds of contacts made by lysine sidechains in the parental constructs (**Table 2** and **Fig. 5**). The larger number of crystal-packing contacts made by the engineered *vs.* native arginine residues could potentially reflect greater sequestration of the native residues in local surface interactions reducing the probability of reaching across a packing interface to make an energetically stabilizing interaction with a neighboring molecule in the crystal lattice. More extensive experimentation will be required to evaluate this possibility and to establish the statistical robustness of the trends documented in **Table 2**, but they nonetheless support the premise underlying our Bulk KR substitution strategy, which was based on the substantially stronger overrepresentation of arginine *vs.* lysine in crystal-packing interfaces in our large-scale analysis of crystal structures previously deposited in the PDB (**Fig. 1**).

**Table 2.**
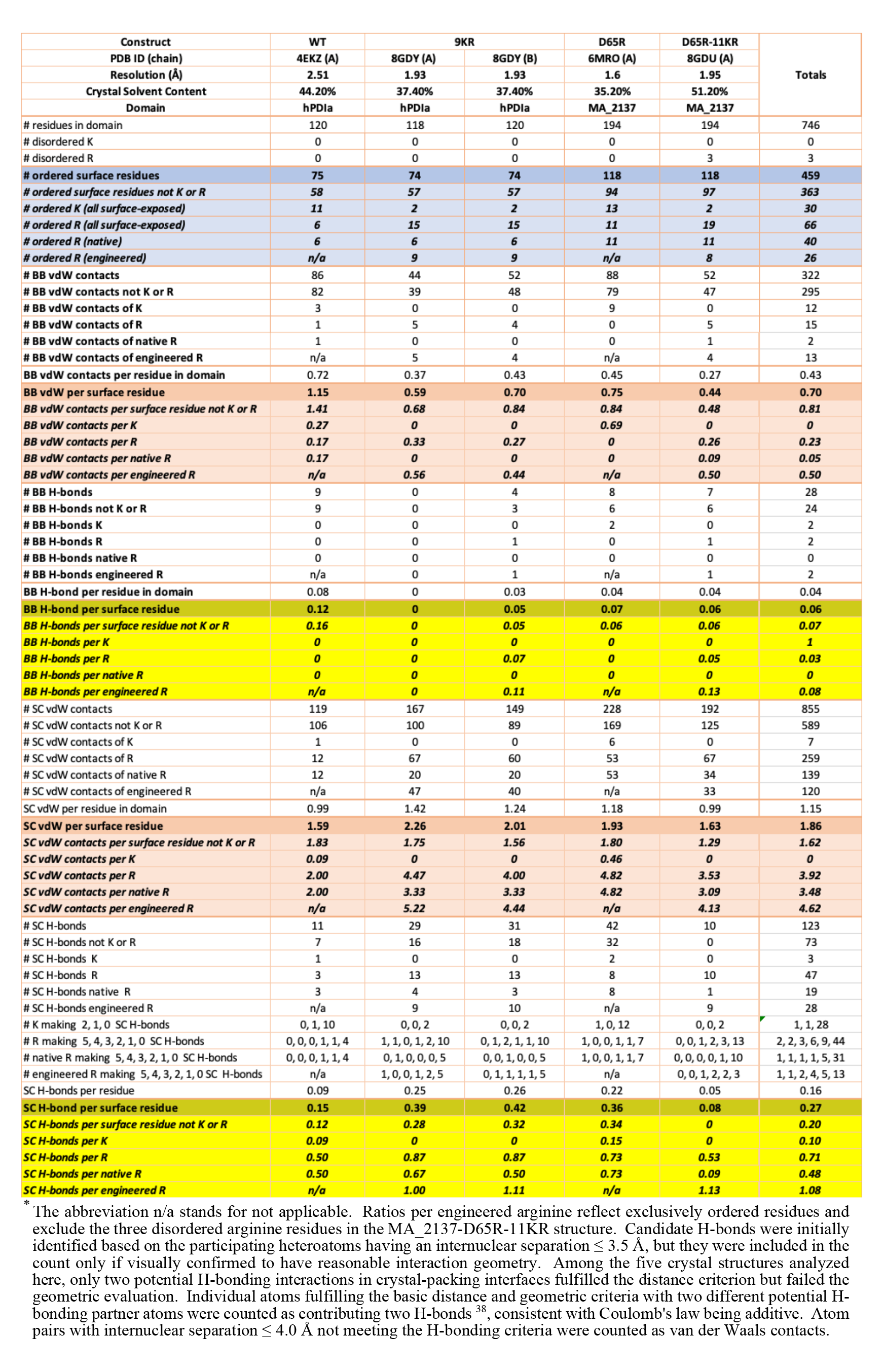
Crystal-packing contacts in reference and Bulk KR protein structures.*

| Construct | WT | 9KR |  | D65R | D65R-11KR | Totals |
| --- | --- | --- | --- | --- | --- | --- |
| PDB ID (chain) | 4EKZ (A) | 8GDY (A) | 8GDY (B) | 6MRO (A) | 8GDU (A) |  |
| Resolution (Å) | 2.51 | 1.93 | 1.93 | 1.6 | 1.95 |  |
| Crystal Solvent Content | 44.20% | 37.40% | 37.40% | 35.20% | 51.20% |  |
| Domain | hPDla | hPDla | hPDla | MA_2137 | MA_2137 |  |
| # residues in domain | 120 | 118 | 120 | 194 | 194 | 746 |
| # disordered K | 0 | 0 | 0 | 0 | 0 | 0 |
| # disordered R | 0 | 0 | 0 | 0 | 3 | 3 |
| # ordered surface residues | 75 | 74 | 74 | 118 | 118 | 459 |
| # ordered surface residues not K or R | 58 | 57 | 57 | 94 | 97 | 363 |
| # ordered K (all surface-exposed) | 11 | 2 | 2 | 13 | 2 | 30 |
| # ordered R (all surface-exposed) | 6 | 15 | 15 | 11 | 19 | 66 |
| # ordered R (native) | 6 | 6 | 6 | 11 | 11 | 40 |
| # ordered R (engineered) | n/a | 9 | 9 | n/a | 8 | 26 |
| # BB vdW contacts | 86 | 44 | 52 | 88 | 52 | 322 |
| # BB vdW contacts not K or R | 82 | 39 | 48 | 79 | 47 | 295 |
| # BB vdW contacts of K | 3 | 0 | 0 | 9 | 0 | 12 |
| # BB vdW contacts of R | 1 | 5 | 4 | 0 | 5 | 15 |
| # BB vdW contacts of native R | 1 | 0 | 0 | 0 | 1 | 2 |
| # BB vdW contacts of engineered R | n/a | 5 | 4 | n/a | 4 | 13 |
| BB vdW contacts per residue in domain | 0.72 | 0.37 | 0.43 | 0.45 | 0.27 | 0.43 |
| BB vdW per surface residue | 1.15 | 0.59 | 0.70 | 0.75 | 0.44 | 0.70 |
| BB vdW contacts per surface residue not K or R | 1.41 | 0.68 | 0.84 | 0.84 | 0.48 | 0.81 |
| BB vdW contacts per K | 0.27 | 0 | 0 | 0.69 | 0 | 0 |
| BB vdW contacts per R | 0.17 | 0.33 | 0.27 | 0 | 0.26 | 0.23 |
| BB vdW contacts per native R | 0.17 | 0 | 0 | 0 | 0.09 | 0.05 |
| BB vdW contacts per engineered R | n/a | 0.56 | 0.44 | n/a | 0.50 | 0.50 |
| # BB H-bonds | 9 | 0 | 4 | 8 | 7 | 28 |
| # BB H-bonds not K or R | 9 | 0 | 3 | 6 | 6 | 24 |
| # BB H-bonds K | 0 | 0 | 0 | 2 | 0 | 2 |
| # BB H-bonds R | 0 | 0 | 1 | 0 | 1 | 2 |
| # BB H-bonds native R | 0 | 0 | 0 | 0 | 0 | 0 |
| # BB H-bonds engineered R | n/a | 0 | 1 | n/a | 1 | 2 |
| BB H-bond per residue in domain | 0.08 | 0 | 0.03 | 0.04 | 0.04 | 0.04 |
| BB H-bond per surface residue | 0.12 | 0 | 0.05 | 0.07 | 0.06 | 0.06 |
| BB H-bonds per surface residue not K or R | 0.16 | 0 | 0.05 | 0.06 | 0.06 | 0.07 |
| BB H-bonds per K | 0 | 0 | 0 | 0 | 0 | 1 |
| BB H-bonds per R | 0 | 0 | 0.07 | 0 | 0.05 | 0.03 |
| BB H-bonds per native R | 0 | 0 | 0 | 0 | 0 | 0 |
| BB H-bonds per engineered R | n/a | 0 | 0.11 | n/a | 0.13 | 0.08 |
| # SC vdW contacts | 119 | 167 | 149 | 228 | 192 | 855 |
| # SC vdW contacts not K or R | 106 | 100 | 89 | 169 | 125 | 589 |
| # SC vdW contacts of K | 1 | 0 | 0 | 6 | 0 | 7 |
| # SC vdW contacts of R | 12 | 67 | 60 | 53 | 67 | 259 |
| # SC vdW contacts of native R | 12 | 20 | 20 | 53 | 34 | 139 |
| # SC vdW contacts of engineered R | n/a | 47 | 40 | n/a | 33 | 120 |
| SC vdW per residue in domain | 0.99 | 1.42 | 1.24 | 1.18 | 0.99 | 1.15 |
| SC vdW per surface residue | 1.59 | 2.26 | 2.01 | 1.93 | 1.63 | 1.86 |
| SC vdW contacts per surface residue not K or R | 1.83 | 1.75 | 1.56 | 1.80 | 1.29 | 1.62 |
| SC vdW contacts per K | 0.09 | 0 | 0 | 0.46 | 0 | 0 |
| SC vdW contacts per R | 2.00 | 4.47 | 4.00 | 4.82 | 3.53 | 3.92 |
| SC vdW contacts per native R | 2.00 | 3.33 | 3.33 | 4.82 | 3.09 | 3.48 |
| SC vdW contacts per engineered R | n/a | 5.22 | 4.44 | n/a | 4.13 | 4.62 |
| # SC H-bonds | 11 | 29 | 31 | 42 | 10 | 123 |
| # SC H-bonds not K or R | 7 | 16 | 18 | 32 | 0 | 73 |
| # SC H-bonds K | 1 | 0 | 0 | 2 | 0 | 3 |
| # SC H-bonds R | 3 | 13 | 13 | 8 | 10 | 47 |
| # SC H-bonds native R | 3 | 4 | 3 | 8 | 1 | 19 |
| # SC H-bonds engineered R | n/a | 9 | 10 | n/a | 9 | 28 |
| # K making 2, 1, 0 SC H-bonds | 0, 1, 10 | 0, 0, 2 | 0, 0, 2 | 1, 0, 12 | 0, 0, 2 | 1, 1, 28 |
| # R making 5, 4, 3, 2, 1, 0 SC H-bonds | 0, 0, 0, 1, 1, 4 | 1, 1, 0, 1, 2, 10 | 0, 1, 2, 1, 1, 10 | 1, 0, 0, 1, 1, 7 | 0, 0, 1, 2, 3, 13 | 2, 2, 3, 6, 9, 44 |
| # native R making 5, 4, 3, 2, 1, 0 SC H-bonds | 0, 0, 0, 1, 1, 4 | 0, 1, 0, 0, 0, 5 | 0, 0, 1, 0, 0, 5 | 1, 0, 0, 1, 1, 7 | 0, 0, 0, 0, 1, 10 | 1, 1, 1, 1, 5, 31 |
| # engineered R making 5, 4, 3, 2, 1, 0 SC H-bonds | n/a | 1, 0, 0, 1, 2, 5 | 0, 1, 1, 1, 1, 5 | n/a | 0, 0, 1, 2, 2, 3 | 1, 1, 2, 4, 5, 13 |
| SC H-bonds per residue | 0.09 | 0.25 | 0.26 | 0.22 | 0.05 | 0.16 |
| SC H-bond per surface residue | 0.15 | 0.39 | 0.42 | 0.36 | 0.08 | 0.27 |
| SC H-bonds per surface residue not K or R | 0.12 | 0.28 | 0.32 | 0.34 | 0 | 0.20 |
| SC H-bonds per K | 0.09 | 0 | 0 | 0.15 | 0 | 0.10 |
| SC H-bonds per R | 0.50 | 0.87 | 0.87 | 0.73 | 0.53 | 0.71 |
| SC H-bonds per native R | 0.50 | 0.67 | 0.50 | 0.73 | 0.09 | 0.48 |
| SC H-bonds per engineered R | n/a | 1.00 | 1.11 | n/a | 1.13 | 1.08 |

\* The abbreviation n/a stands for not applicable. Ratios per engineered arginine reflect exclusively ordered residues and exclude the three disordered arginine residues in the MA\_2137-D65R-11KR structure. Candidate H-bonds were initially identified based on the participating heteroatoms having an internuclear separation $\leq 3.5$ Å, but they were included in the count only if visually confirmed to have reasonable interaction geometry. Among the five crystal structures analyzed here, only two potential H-bonding interactions in crystal-packing interfaces fulfilled the distance criterion but failed the geometric evaluation. Individual atoms fulfilling the basic distance and geometric criteria with two different potential H-bonding partner atoms were counted as contributing two H-bonds<sup>38</sup>, consistent with Coulomb's law being additive. Atom pairs with internuclear separation $\leq 4.0$ Å not meeting the H-bonding criteria were counted as van der Waals contacts.

On average, the well-ordered engineered arginine sidechains in our Bulk KR structures make 1.08 H-bonds each to a neighboring protein molecule in the crystal lattice, compared to 0.48 each for the native arginines (**Table 2**). In comparison, the well-ordered lysine sidechains make an average of 0.10 H-bonds each to a neighboring protein molecule in the native structures and none in our Bulk KR structures. These results support our hypothesis for the physicochemical basis of the greater overrepresentation of arginine compared to lysine in crystal-packing interfaces in the PDB, which is that that guanidino group in arginine is substantially more efficacious than the primary amine group in lysine in mediating energetically stabilizing H-bonds in the relevant stereochemical contexts (**Fig. 5**).

The number of van der Waals contacts per ordered sidechain follow a similar trend. Engineered and native arginines, respectively, make on average 4.62 and 3.43 contacts each, while lysines in the native structures make on average 0.28 contacts each, and lysines in the engineered structures making none (**Table 2**). The greater number of intermolecular van der Waals contacts made by the arginine sidechains could potentially be influenced by their greater H-bonding propensity leading to more frequent occurrence in crystal-packing interfaces, but additional research will be required to determine the relative energetic contributions of their van der Waals *vs.* H-bonding interactions to lattice stabilization. Notably, the number of backbone H-bonding and van der Waals interactions make by arginine *vs.* lysine residues in our reference and engineered structures do not show any clear trends (**Table 2**).

### Influence of Bulk KR mutations on protein solubility in PEG3350 solutions

Thermodynamic solubility assays using polyethylene glycol 3350 (PEG3350) to induce protein precipitation ^64–66^ assess the relative free energy of the hydrated state of individual protein molecules compared to the most favorable self-associated state under conditions of constant ionic strength but reduced water activity (effective concentration). In practice, these assays monitor optical density at 280 nm in the supernatant of solutions containing different concentrations of protein in the presence of increasing concentrations of PEG3350 after centrifugation to remove large particulate molecular assemblies, so they effectively measure the equilibrium concentration of protein that remains soluble as water activity is reduced. The observed results depend intrinsically on the free energy of the self-associated state, which varies significantly for different proteins in different solvent environments and can include crystalline phases and liquid-liquid phase separated (LLPS) phases in addition to heterogeneous amorphously precipitated phases. This factor can complicate interpretation of thermodynamic solubility assays, but they nonetheless provide insight into physicochemical properties that ultimately control protein crystallization behavior.

PEG3350 precipitation assays on WT hPDIa and the 9KR mutant show no significant difference in their behavior in PEG3350 precipitation assays (**Fig. ED3a**), indicating that the Bulk KR substitutions in this protein domain do not alter its thermodynamic solubility under these conditions even though they enable crystallization and high-resolution structure determination of a domain that does not crystallize at all with its native sequence. Even when harboring the 9KR mutations, hPDIa crystallizes only under a very small fraction (0.6%) of the solution conditions explored in high-throughput crystallization screening (**Fig. 4**) while showing amorphous precipitation in many of them (data not shown). Therefore, our thermodynamic solubility assays on the hPDIa constructs are likely measuring the free energy of the hydrated state of individual protein molecules compared to amorphously precipitated phases, and they demonstrate that the physicochemical properties controlling the formation of such phases are likely different from those controlling protein crystallization behavior.

PEG3350 precipitation assays on our MA_2137 constructs demonstrate more complex phase behavior (**Fig. ED3b-c**) likely reflecting different physical forms of self-association under assay conditions. Notably, the WT and D65R mutant could not be precipitated by the highest 35% (v/v) concentration of PEG3350 that was assayed (**Fig. ED3b**). These protein constructs instead showed some tendency to exhibit a small increase in optical density at low PEG3350 concentration, likely reflecting light scattering due some form of protein self-association in a low-density state that does not sediment during low-speed centrifugation. During crystallization screening, these constructs showed clear evidence of liquid-liquid phase separation LLPS without any apparent amorphous precipitation in many reaction conditions (**Fig. ED4**). Therefore, the inability to precipitate these constructs at high PEG3350 likely concentration likely reflects LLPS being energetically more favorable for this protein under conditions of low water activity than amorphous precipitation. Our crystal structures of MA_2137 constructs show clear and well-ordered electron density for every residue in this 202-residue protein except for the C-terminal hexahistidine tag that was added to enable purification using NiNTA affinity chromatography and a 12-residue internal loop that is disordered in the MA_2137-D65R-11KR structure, although well ordered by Ca^++^ ions from the mother liquor in the structure of the parental MA_2137-D65R construct (**Fig. ED2b**). Furthermore, our CD thermal melting data demonstrate that the protein is very stably folded (**Fig. 3**). Therefore, our solubility data (**Fig. ED3b**) combined with our crystallization screening data (**Fig. ED4**) suggest that MA_2137 undergoes LLPS in an essentially fully folded conformational state.

In contrast to the behavior of the WT and the D65R constructs, the 5KR and 11KR constructs of MA_2137 show precipitation at the highest PEG3350 concentrations used in our solubility assays, with the 11KR construct showing stronger precipitation than the 5KR construct (**Fig. ED3c**). These results indicate the free energy of these MA_2137 constructs is lower in the precipitated state than in the LLPS state under conditions of very low water activity, reflecting a reduction in thermodynamic solubility. However, these constructs both crystallize extremely promiscuously, with the 5KR and 11KR constructs yielding crystallization hits in ∼8% and ∼15% of screened conditions, respectively (**Fig. 4**). These results raise the possibility that the precipitate formed by the 5KR and 11KR constructs at very high PEG3350 concentration could be in a microcrystalline state rather than amorphously precipitated state due to the high efficacy of the Bulk KR mutations in promoting crystallization. Further research will be needed to determine whether the reduced solubility of the 5KR and 11KR constructs reflects stabilization of crystalline states or amorphously precipitated states of MA_2137-D65R.

## Discussion

The results presented in this paper demonstrate the efficacy of a new method for probabilistic engineering of protein surface properties to enhance crystallization propensity based on substitution of multiple lysine (K) residues with arginine (R). The rationale behind this “Bulk KR” substitution method is that lysine and arginine have very similar physicochemical properties, but arginine shows substantially higher overrepresentation than lysine in a large-scale computational analysis we performed of crystal structures deposited in the PDB (**Fig. 1**). We have developed software to rank lysine sites for substitution based on the redundancy-corrected count of KR substitutions observed in homologous proteins with the highest level of sequence identity (**Fig. 2**), based on the rationale that biological evolution selects against destabilizing and function-impairing mutations. We demonstrate that mutations selected this way are only minimally destabilizing (**Fig. 3** and **Table 1**) and significantly enhance crystallization propensity for two of three test proteins (**Fig. 4**). The crystals yielded by our Bulk KR method diffract strongly and enabled efficient determination of a 1.9 Å crystal structure (**Table ED1**) for the hPDIa protein domain that does not crystallize at all with its native sequence (**Fig. 4** and **Table 1**). Our crystal structures of Bulk KR substituted proteins show no significant conformational or stereochemical differences *vs.* reference proteins (**Fig. ED2**), and the engineered arginine residues, like the native ones, making both van der Waals contacts and H-bonds in crystal-packing interfaces at substantially higher frequencies than either lysine residues or other residues (**Table 2**). These crystal structures were produced by the Bulk KR constructs harboring the highest number of substitutions, which were the only constructs for which any diffraction data were measured. These results support the efficacy of a streamlined pipeline for crystal structure determination in which solubility is tested for a set of constructs with an increasing number of KR mutations but purification and crystallization screening is only performed on the construct harboring the largest number of mutations. In summary, the biophysical results presented in this paper support bulk KR substitution being a rational and effective probabilistic strategy to engineer protein surface properties to enhance protein crystallization propensity.

Our Bulk KR method focuses on large-scale modification of protein surface properties while preserving physicochemical properties at every site. KR mutations have been explored in the past both for their ability to modulate protein stability and crystallization propensity. Bulk KR substitution in GFP was shown to greatly reduce the amount of soluble protein expressed *in vivo* in *E. coli* and also to reduce the fluorescence level of the protein that could be purified, although the mutations slowed the rate of unfolding by chemical denaturants ^52^. However, this study only examined 14KR an 19KR mutations at sites selected based on diffuse criteria. Our studies show that KR mutations at 25 sites selected based on the frequency of substitution observed in homologs shows on average a 0.54 °C reduction in T_m_ per KR mutation (**Fig. 3** and **Table 1**).

The previous studies of the influence of KR mutations on crystallization propensity were motivated by an earlier computational study using different stereochemical and statistical normalization methods to analyze the amino acids making crystal-packing interactions in a small set of 233 protein crystal structures ^67^. This analysis produced very different conclusions when comparing results for all 20 amino acids (**Fig. ED5**), with the most salient difference being the conclusion that lysine, glutamate, and tryptophan are all disfavored in crystal-packing contacts, directly contradictory to the results of our computational analysis of 87,684 crystal structures (**Fig. 1**). Supporting the inaccuracy of this earlier conclusion and the validity of our analysis, we have successfully used introduction of both lysine and glutamate residues to improve protein crystallization (manuscript in preparation). The earlier computational analysis did conclude that arginine is favored in crystal-packing contacts, leading the authors to suggest that KR substitutions could improve protein crystallization propensity, but they did not perform any experiments testing this proposal. Two different groups subsequently tested this proposal using a small number of KR substitutions ^24, 27^. Both concluded that KR substitution show limited efficacy in improving protein crystallization, but one of them was not able to obtain a diffracting crystal using a series of single-site KR substitutions ^27^. The other group was able to obtain a crystal structure, but their overall conclusion was that lysine-to-alanine (KA) substitutions shows substantially greater efficacy in improving protein crystallization properties than KR substitutions^24^. Unfortunately, the KA method has had minimal impact improving protein crystallization based on structures submitted to the PDB during the ∼20 years since it was proposed. One contributor to the lack of successful application of the KA method may be that the grossly different physicochemical properties of alanine compared to lysine can produce significant protein destabilization, which impedes introduction of multiple KA mutations. However, our computational analyses raise additional questions about the KA method because they indicate that alanine is underrepresented in crystal-packing interfaces while lysine is overrepresented (**Fig. 1**).

The conceptual foundation of our crystallization engineering methods is fundamentally different from that of these earlier studies. Our method focuses on probabilistic reengineering of protein surface properties via substitution of multiple amino acids with similar physicochemical properties but different propensity to make crystal-packing interactions based on large-scale computational analyses of previously determined crystal structures. The data presented in this paper support high efficacy for the method including importantly when applied in a streamlined fashion in which a series of mutant protein variants with increasing numbers of physiochemically conservative crystallization-enhancing mutations are evaluated for soluble protein expression and only the construct with greatest number of crystallization-enhancing is purified and subjected to crystallization screening (**Figs. 4-5**, **Tables 1-2**, and **Table ED1**).

Given the success of the Bulk KR method in improving protein crystallization behavior, our computational analysis of crystal-packing interactions in the PDB (**Fig. 1**) suggests several related strategies with promise to improve crystallization behavior based on the same conceptual approach. Aspartate and glutamate frequently substitute for one another in the course of evolution^68–70^ due to their very similar physicochemical properties, but glutamate shows over 2-fold higher overrepresentation in crystal-packing interfaces (**Fig. 1**). A similar trend relative to crystal packing interactions is observed for asparagine and glutamine, which also have very similar physicochemical properties. These observations suggest bulk aspartate-to-glutamate (DE) and asparagine-to-glutamine (NQ) substitutions are also likely to improve crystallization propensity.

In the case of these substitutions, the higher entropy of the sidechain with greater crystal-packing propensity will tend to thermodynamically oppose immobilization in a crystal-packing interface, while this factor does not apply to bulk lysine-to-arginine substitution due to the very similar entropy of these sidechains. However, our computational analysis of crystal-packing interactions in the PDB shows that high-entropy sidechains mediating crystal-packing interactions tend to participate in salt-bridging and H-bonding interactions with nearby residues in the primary sequence, especially at ±3 and ±4 positions in α-helices and ±2 positions in β-strands ^36–38^ (manuscript in preparation). These interactions likely reduce the entropy of the sidechains in the isolated protein molecules, which will reduce or eliminate entropy loss due to immobilization in a crystal-packing interface. Therefore, bulk DE and NQ substitution seems likely to be most effective when residues with potential salt-bridging or H-bonding partners at ±3 and ±4 positions in α-helices or ±2 positions in β-strands are prioritized for substitution.

Future research will be required to assess the efficacy of these alternative bulk substitution methods and to establish the most efficient paradigm for combing KR, DE, and NQ mutations to maximize crystallization hit rate and crystal quality while minimizing the number of constructs needed to obtain a diversity of different crystal forms and a high-resolution crystal structure. Nonetheless, our bulk substitution methods focused on large-scale probabilistic remodeling of protein surface properties to enhance crystallization propensity already show significant efficacy for rational engineering of proteins to improve their crystallization properties.

## Acknowledgements

This work was supported by a grant from the US NIH-NIGMS to JFH (GM127883). We thank Accendro Inc. for assistance with web programming and G.T. Montelione, R. Xiao, and the other members of the Northeast Structural Genomics Consortium for long-term collaboration and advice.

## Author contributions

Conceptualization: JFH. Methodology: NEB, SS, SKH, WNP, HSH, & JFH. Investigation: NEB, BJL, SS, MN, & KHW. Visualization: NEB &JFH. Funding acquisition: JFH. Project administration: JFH. Supervision: JFH. Writing: NEB & JFH.

## Conflict of interest

JFH is a member of the Scientific Advisory Board of Nexomics Biosciences. The other authors declare no competing financial interests.

*Note: The **Supplementary Information** for this manuscript contains 3 figures and 1 table*.

## Methods

### GPU acceleration of sequence identity calculation

Our Python program accelerates calculation of the absolute percent identity between two sequences by parallelizing site-by-site comparison on a Graphical Processing Unit (GPU) chip using a custom reduction kernel ^71^ written using the CuPy library ^72^. In brief, each pair of aligned amino acids in two sequences is sent to a separate GPU core. If the 1-character amino acid codes in both sequences match each other and neither is empty due to a gap in the sequence alignment, that core stores a value of 1 in its register, which is otherwise set to zero. The values in the registers for all sites are then summed using parallel reduction to count the number of identical amino acids in the sequence. Details can be found in the code at https://github.com/huntmolecularbiophysicslab/pxengineering.

### Prioritization of mutation sites based on redundancy-corrected counts of KR mutations observed in homologous proteins

In order to compensate for outright redundancy as well as inhomogeneous phylogenetic sampling in sequence databases, our software performs two redundancy-compensation calculations on the set of sequences in each percent-identity bin having an arginine substitution at a specific lysine site in the target protein. Both calculations use the same heuristic estimate for the probability of evolutionary resampling at an aligned site between two sequences *i* and *j* with an overall fraction of *f_id_* identical residues at all aligned sites:

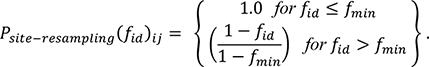

We assume *f_min_* = 0.3. One calculation gives a redundancy-reduced estimate *C_R_* of arginine counts using the following formula in which the summation is performed over all unique pairs of the *N* sequences in the bin having an arginine substitution at one lysine site in the target protein:

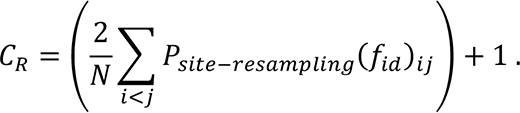

This calculation is extremely rapid and includes all homologs but is only rigorously accurate in the limiting cases of full redundancy or full independence. The second calculation gives a rigorous estimate of the expectation value for the number of independent observations of arginine based on combinations of the heuristic probabilities of being resampled or not being resampled for all unique pairs among the seven most diverged sequences in each bin that have arginine at a specific lysine site. These sequences are identified by taking the single sequence with the lowest percent identity to the target sequence and then progressively adding the sequence with the lowest average percent identity to those already selected. The details of the implementation can be found in the code at https://github.com/huntmolecularbiophysicslab/pxengineering.

### Protein expression and purification

Protein coding sequences harboring Bulk KR Substitutions were synthesized (Twist Bioscience, South San Francisco, CA) with a short C-terminal hexahistidine tag (LEHHHHHH), cloned into the T7 expression plasmid pET21_NESG (https://dnasu.org/DNASU/GetCloneDetail.do?cloneid=336944), and then transformed into BL21(DE3) Rosetta *E. coli* cells (MilliporeSigma, Burlington, MA). Protein expression was induced with 1mM IPTG for 4 hours at 30°C in Terrific Broth ^73^. Cells were pelleted by centrifugation at 4,000 rpm for 25 min at 4°C and resuspended on ice in 10 mM imidazole, 300 mM NaCl, 1 mM TCEP, 5% (w/v) glycerol, 50 mM NaH_2_PO_4_, pH 7.5, prior to cell lysis by probe sonication. The supernatant following 15,000 rpm centrifugation at 4°C was mixed with Ni-NTA resin and incubated at 4 °C for 1 hour. The mixture was then transferred into a column and washed with the same buffer containing a higher 100 mM imidazole concentration prior to elution of the protein in 6 ml of the same buffer containing 250 mM imidazole. A 1mL aliquot of eluted protein was concentrated to 500 µl using an Amicon 10 kDa centrifugal filter (MilliporeSigma, Burlington, MA) and loaded via a 1 ml loop for onto a Superdex 200 Increase 10/300 gl gel filtration column equilibrated in 100 mM NaCl, 10 mM DTT, 10 mM Tris-Cl, pH 7.5. Protein-containing fractions were concentrated to ∼15 mg/ml based on *a priori* sequence-based extinction coefficients ^74^ and OD_280nm_ values measured using a Nanodrop spectrophotometer (ThermoFisher, Waltham, MA), and the concentrated protein was immediately flash-frozen in aliquots in liquid nitrogen prior to storage at −80 °C pending use.

### Thermal stability assays using CD spectroscopy

An Applied Photophysics (Leatherhead, UK) Chirascan V100 spectropolarimeter with a Peltier-jacketed cell holder was used to collect circular dichroism (CD) spectra spanning 200-250 nm serially during a 3 °C/min thermal ramp nominally running from 10-84 °C. Data were measured from protein samples in a 0.5 mm quartz cuvette in 1 nm increments using a 0.25 integration time per point and a 1 nm bandwidth, corresponding to 35 sec per spectrum. Measurements of the actual cell temperature during the experiment, which were used for data display and analysis, indicated the actual range of the temperature ramp was from ∼11-78 °C for all samples. Protein samples were diluted to 2 mg/ml using gel filtration buffer lacking DTT (*i.e.*, 100 mM NaCl, 10 mM Tris-Cl, pH 7.5) with the addition of 1 mM SAH for the MA_237 constructs. Global curve fitting of spectral data during the thermal ramp was performed from 215-230 nm using the program GLOBAL3 (Applied Photophysics) using double linear baseline correction (*i.e.*, before and after the observed transitions) to extract the thermodynamic parameters and melting temperatures (**Table 1** and **Fig. 3**). Each dataset was analyzed using the smallest number of transitions showing approximately random directions for the residuals for adjacent points in the CD *vs.* measured temperature plane.

### Solubility Assays

Protein stock solutions were diluted to working concentration in the same buffer used for gel-filtration chromatography containing different weight/volume concentrations of PEG3350. Following a 60 minute incubation at room temperature, the samples were spun for 10 min at 14,000 RPM in a microfuge to pellet particulates, and the concentration of protein in the supernatant was measured using the optical density at 280 nm measured in a Nanodrop spectrophotometer (ThermoFisher) based on the *a priori* extinction coefficient ^74^. The centrifugation and measurement of concentration in the supernatant were repeated 24 hours later to ensure equilibrium had been reached.

### Protein crystallization

A subset of the crystal hits for hPDIa-9KR and MA_2137-D65R-11KR observed in the 1536-well screen conducted at the High-Throughput Crystallization Screening Center ^53–58^ at the Hauptman-Woodward Medical Research Institute (https://hwi.buffalo.edu/high-throughput-crystallization-center/) using 15 mg/ml protein stock concentrations were initially reproduced using the microbatch-under-oil method at 4 ℃ and 18 ℃ and subsequently optimized by seeding. The hPDIa-9KR stock was mixed at a 2:1 volume ration with a crystallization reagent comprising 24% (w/v) PEG 20k, 0.1 M potassium thiocyanate, 0.1 M MES, pH 6. The MA_2137- D65R-11KR stock was mixed at a 1:1 volume ration with a crystallization reagent comprising 30% (w/v) PEG 1k, 0.1 M HEPES, pH 7.5. All crystals were transferred into a similar crystallization solution supplemented with 20% (v/v) ethylene glycol prior to mounting and flash-freezing in liquid nitrogen. The WT and 5KR RNaseH constructs were screened in the same manner also using a protein stock concentration of 15 mg/ml but showed pervasive amorphous precipitation without yielding any crystallization hits, suggesting screening may have been conducted at too high a protein concentration.

### Crystal structure determination and refinement

X-ray diffraction data were collected from single crystals of PDIa-9M and MA_2137-D65R-11 using, respectively, the NE-CAT 24-ID-E and 24-ID-C beam lines at the Advanced Photon Source (**Table ED1**). The images were processed and scaled using XDS ^75–78^. The structure of hPDIa-9KR was solved by molecular replacement using the program MOLREP ^79^ employing a search model comprising the first domain in the crystal structure of full-length hPDI (PDB id 4EKZ). The structure of MA_2137-D65R-11KR was solved using the same methods employing the structure of MA_2137-D65R (PDB id 6MRO) as the search model. Both structures were refined (**Table ED1**) using PHENIX ^4^ in conjunction with manual rebuilding in XtalView ^80^ and COOT ^81^.

