## Extended Data for "Systematic enhancement of protein crystallization efficiency by bulk lysine-to-arginine (KR) substitution"

*Extended Data*
*for*

### Systematic enhancement of protein crystallization efficiency by bulk lysine-to-arginine (KR) substitution

**Nooriel E. Banayan [1†], Blaine J. Loughlin [1†], Shikha Singh [1], Farhad Forouhar [1], Guanqi Lu [1], Kam-Ho Wong [1§], Matthew Neky [1], Henry S. Hunt [2], Larry B. Bateman Jr. [3], Angel Tamez [3], Samuel K. Handelman [1§], W. Nicholson Price [1§], & John F. Hunt [1*]**

[1] Department of Biological Sciences, 702A Sherman Fairchild Center, MC2434, Columbia University, New York, NY, 10027, USA;

[2] Physics Department, Stanford University, Stanford, CA 94305, USA; and

[3] Accendero Software, P.O. Box 2826, Idaho Falls, ID 83404, USA.

[†] These authors contributed equally to the work reported in this paper.

[*] To whom correspondence may be addressed:

[§] Current addresses:
SKH, 307 E. Merrill St, Indianapolis, IN 46225; MN, Mailman School of Public Health, Columbia University, 722 W. 168th Street, New York, NY 10032; WNP, University of Michigan Law School, 625 S. State St, Ann Arbor, MI 48109; KHW, 401 N Middletown Rd, Pearl River, NY 10965.

#### Bulk K-to-R substitution enhances crystallization

**Keywords: protein crystallization / protein thermodynamics / protein engineering / homology analysis / circular dichroism spectroscopy / protein solubility / x-ray crystallography**

### Extended Data Table 1

***Data collection & refinement statistics for Bulk-KR crystal structures.***

| **Protein** | **hPDIa-9KR** | **MA_2137-D65R-11KR** |
|---|---|---|
| **Crystal parameters** | | |
| Space group | $P2_12_12_1$ | $P4_22_12$ |
| Cell dimensions *a*, *b*, *c* (Å) | 58.7, 61.3, 68.6 | 109.9, 109.9, 39.1 |
| Cell angles $\alpha, \beta, \gamma$ (°) | 90, 90, 90 | 90, 90, 90 |
| **Data Collection** | | |
| Resolution (Å) | 68.6-1.89 (1.93-1.89)* | 110-1.91 (1.95-1.92)* |
| $R_{merge}$ (%) | 0.24 (2.15) | 0.135 (2.828) |
| I/σI | 7 (0.9) | 20.4 (0.9) |
| Completeness (%) | 98.8 (82.2) | 99.8 (97.7) |
| Redundancy | 12.8 (10.4) | 25.4 (17.3) |
| CC1/2 | 0.99 (0.46) | 1.00 (0.49) |
| **Model contents** | | |
| Protein residues | A20-137, B18-137 | A1-140,153-194 |
| # protein atoms | 1,897 | 1,464 |
| Ligands | 2 SCN, 1 EDO | 1 SAH |
| # ligand atoms | 10 | 26 |
| # waters | 144 | 82 |
| **Refinement** | | |
| Resolution (Å) | 45.70-2.05 (2.12-2.05)* | 77.75-2.01 (2.07-2.01)* |
| No. reflections | 16,013 (1,574) | 16,541 (1,642) |
| $R_{work}$ / $R_{free}$ (%) | 19.5 (28.8) / 23.2 (34.1) | 18.2 (20.9) / 22.1 (24.4) |
| **Ramachandran Plot (%)** | | |
| Outliers | 0.00 | 0.00 |
| Allowed | 2.99 | 2.25 |
| Favored | 97.01 | 97.75 |
| **B-factors** | | |
| Protein | 34.6 | 43.2 |
| Ligand/ion | 37.9 | 31.8 |
| Water | 39.5 | 53.3 |
| **r.m.s deviations** | | |
| Bond lengths (Å) | 0.007 | 0.007 |
| Bond angles (°) | 0.841 | 0.935 |
| **PDB ID** | **8GDY** | **8GDU** |

* Values in parentheses represent statistics for the highest resolution shell.

### Extended Data Figure 1.

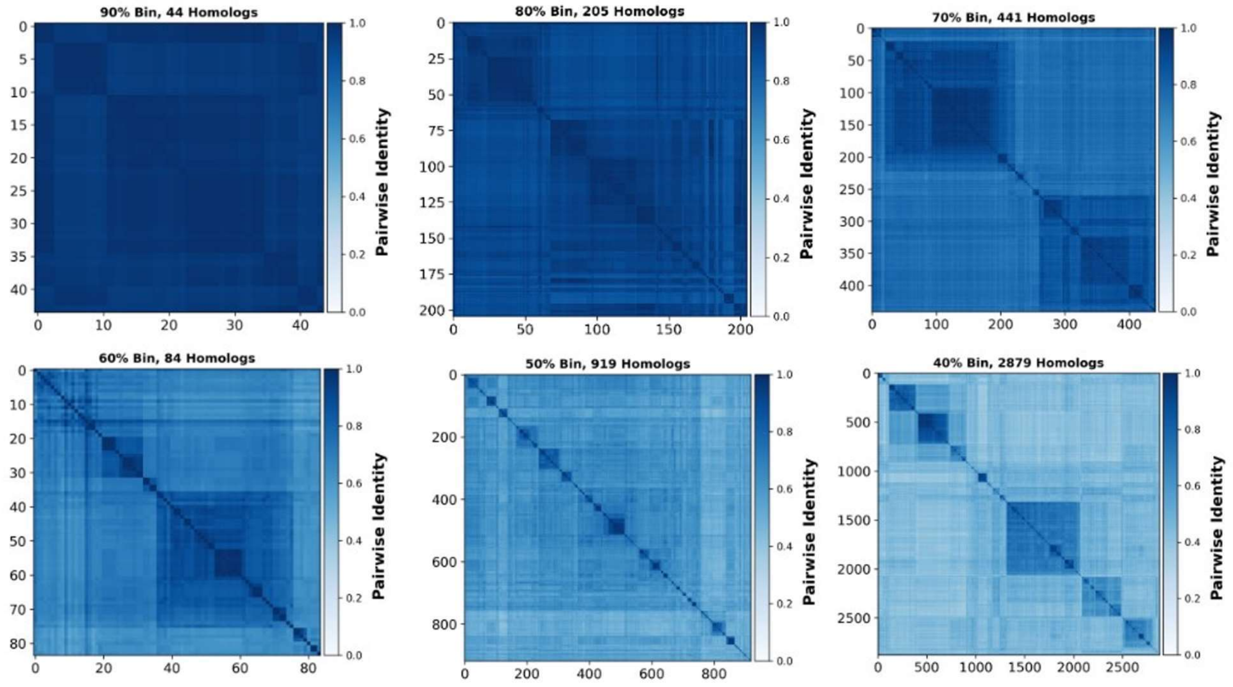


**Figure ED1.** ***Pairwise percent sequence identity distributions between homologs in bins spanning 10% ranges of sequence identity to hPDIa.***

### Extended Data Figure 2.

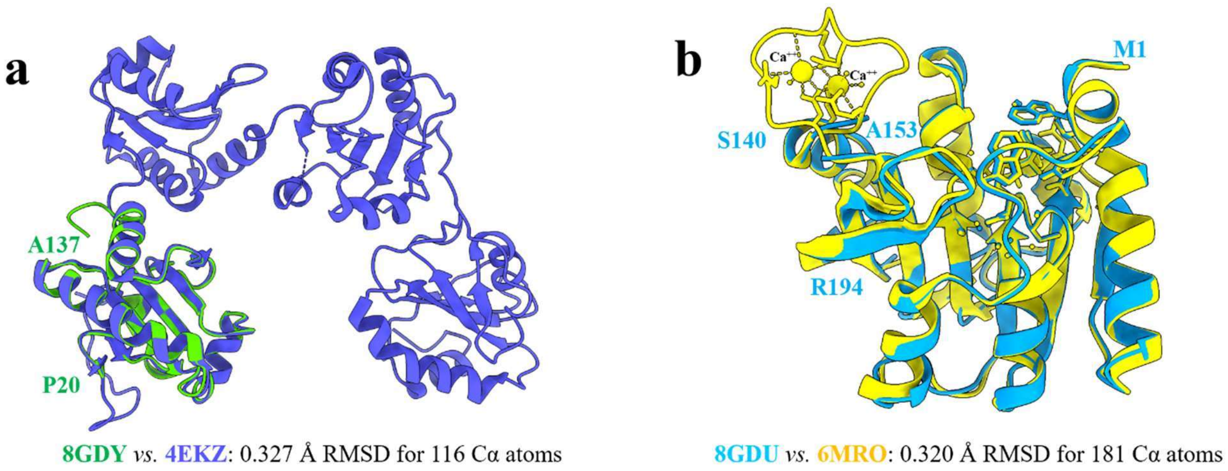


**Figure ED2.** ***Crystal structures of Bulk-KR-substituted protein constructs show no significant conformational or stereochemical changes compared to reference structures.*** **(a)** Comparison of our crystal structure of hPDIa-9KR (**Fig. 5b** and **Table ED1**) to an earlier structure of the same domain as part of a much larger multi-domain hPDI(abb'xa') construct (PDB ID 4EKZ [41]). **(b)** Comparison of our crystal structures of MA_2137-D65R-11KR (**Fig. 5d** and **Table ED1**) and MA_2137-D65R (PDB ID 6MRO). The loop at residues 141-152 is ordered by a high concentration of $Ca^{++}$ present in the mother liquor for the latter structure, while the was no $Ca^{++}$ present in the mother liquor for former structure. Least-squares alignments were performed in PyMOL (www.pymol.org) and ChimeraX [84] using the default algorithms.

### Extended Data Figure 3.

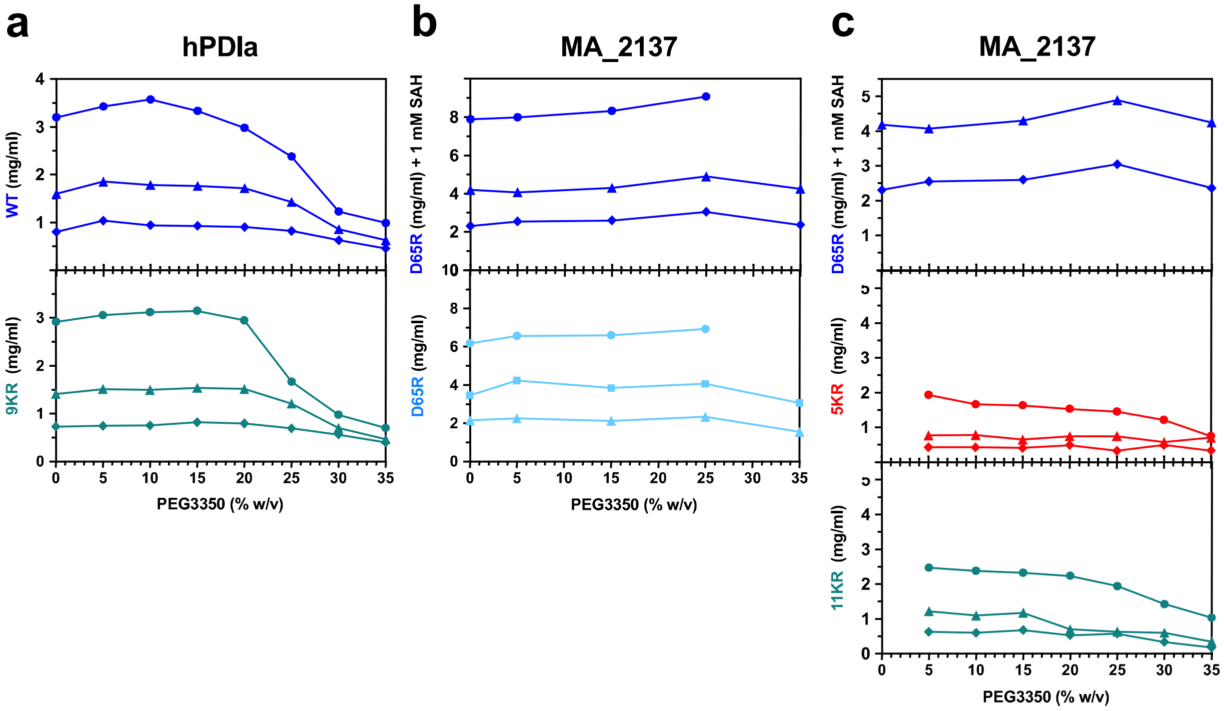


**Figure ED3.** ***Solubility assays on selected hPDIa and MA_2137 constructs.*** Assays were conducted at room temperature. The lower graph in the center column shows data collected without S-adenosylhomocysteine (SAH) in the buffer, which is a product of the methyltransferase reaction catalyzed by MA_2137, while all of the other solubility assays on this protein were conducted in the presence of 1 mM SAH. The top graphs in both MA_2137 columns show the same data, which are duplicated but matched in scale to the graphs beneath them in order to provide a visual reference.

### Extended Data Figure 4.

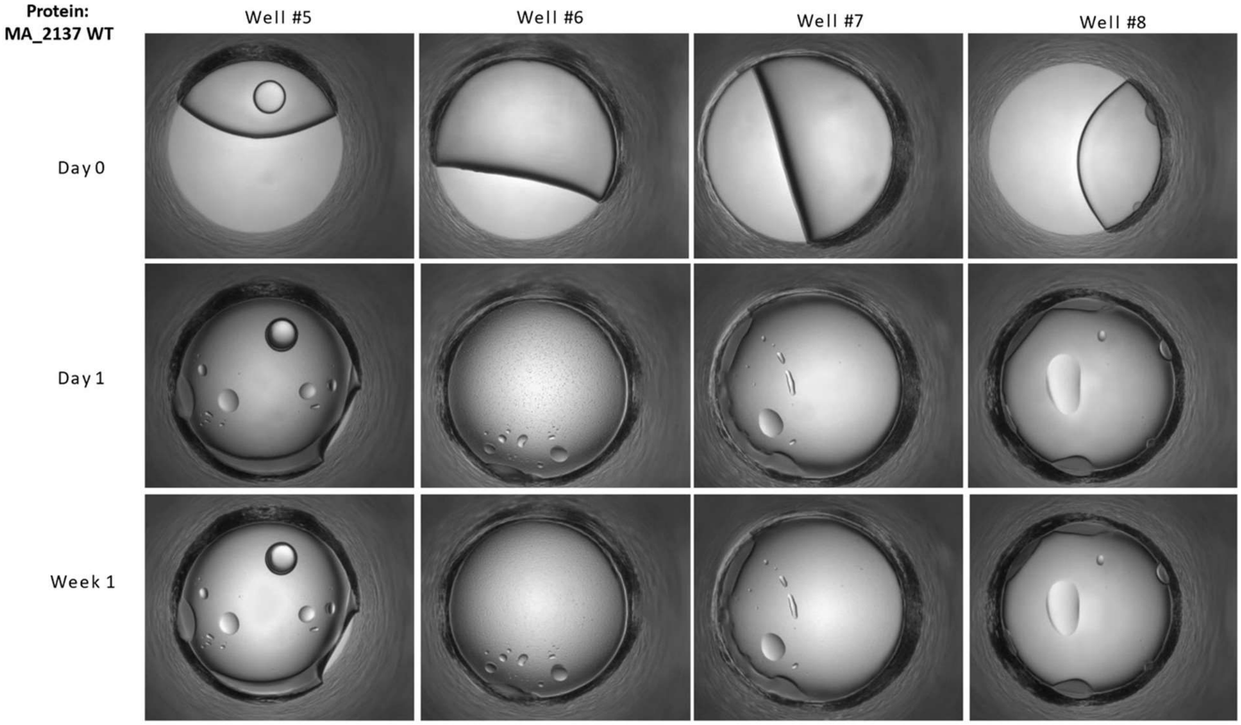


**Figure ED4.** ***Optical micrographs showing time evolution of representative MA_2137-D65R crystallization screening reactions showing evidence of liquid-liquid phase separation.*** Representative micrographs from the HWI generation #19 1,536 well crystal screen (https://hwi.buffalo.edu/crystallization-cocktails/) show evidence of liquid-liquid phase separation for MA_2137-D65R. Other than a tag and 12-residue internal loop, likely to be a calcium binding site (**Fig. ED2b**), this protein is ordered. The cocktail recipes are as follows for the individual wells:

#5: 0.1 M $Na_2S_2O_3$, 0.1 M Tris, pH 8, 20% (w/v) PEG 4000;

#6: (HR SaltRx Screen HT A12) 3.2 M NaCl, 0.1 M NaOAc, pH 4.6;

#7: (HR SaltRx Screen HT D4) 6.0 M $NH_4NO_3$, 0.1M NaOAc, pH 4.6;

#8: 0.1 M LiBr, 0.1 M Tris, pH 8, 20% (w/v) PEG 4000.

#### Extended Data Figure 5.

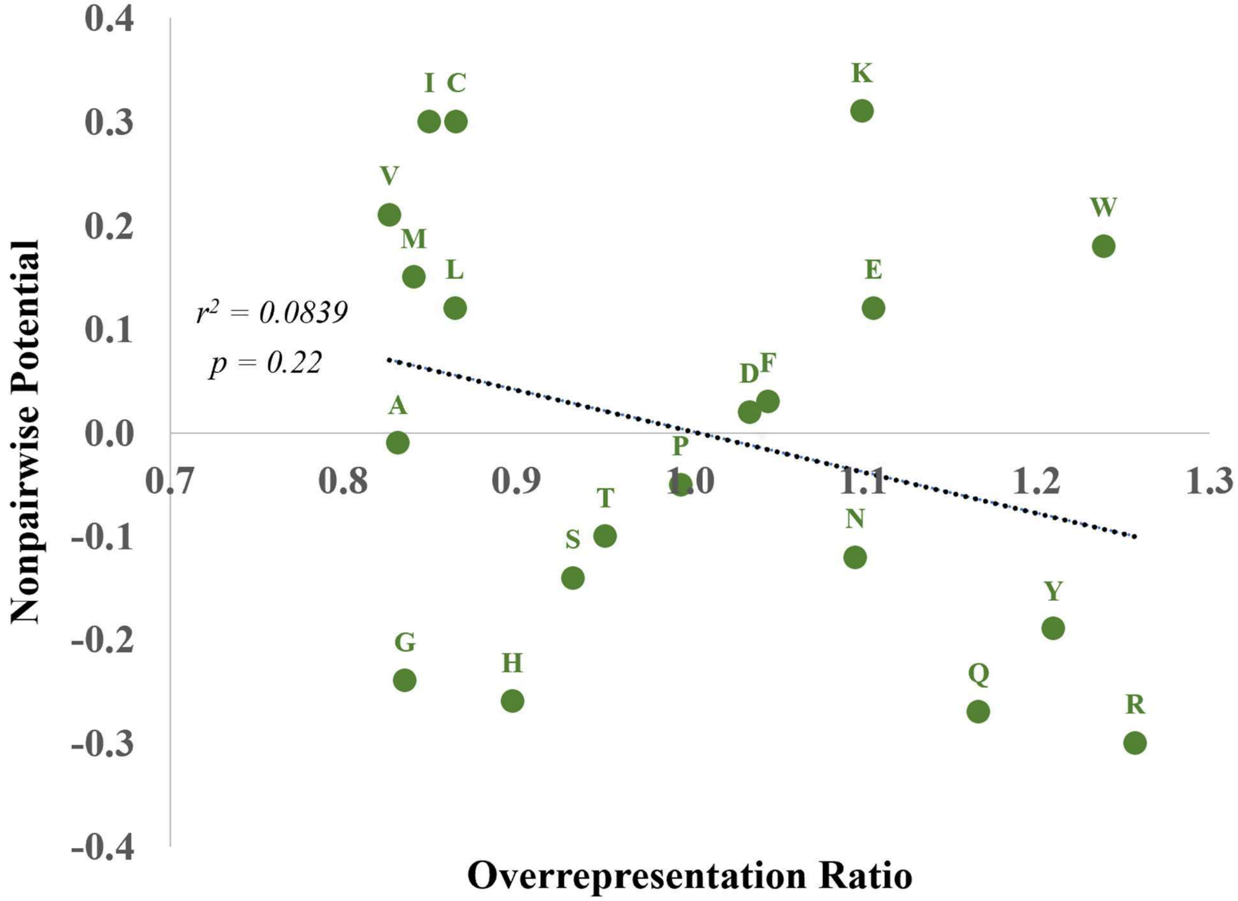


**Figure ED5.** ***Comparison of overrepresentation ratios of amino acids in crystal-packing interfaces in our analysis of 87,684 crystal structures [28] (Fig. 1) to an earlier analysis of contact potentials in crystal-packing interfaces in 233 crystal structures [67].***
